# Common risk variants in *AHI1* are associated with childhood steroid-sensitive nephrotic syndrome

**DOI:** 10.1101/2022.10.05.510923

**Authors:** Mallory L Downie, Sanjana Gupta, Catalin Voinescu, Adam P Levine, Omid Sadeghi-Alavijeh, Stephanie Dufek-Kamperis, Jingjing Cao, Martin Christian, Jameela A Kari, Shenal Thalgahagoda, Randula Ranawaka, Asiri Abeyagunawardena, Rasheed Gbadegesin, Rulan Parekh, Robert Kleta, Detlef Bockenhauer, Horia C Stanescu, Daniel P Gale

## Abstract

**Background:** Steroid-sensitive nephrotic syndrome (SSNS) is the most common form of kidney disease in children worldwide. Genome-wide association studies (GWAS) have demonstrated association of SSNS with genetic variation at *HLA-DQ/DR* and have identified several non-*HLA* loci that aid in further understanding of disease pathophysiology. We sought to identify additional genetic loci associated with SSNS in children of Sri Lankan and European ancestry.

**Methods:** We conducted a GWAS in a cohort of Sri Lankan individuals comprising 420 pediatric patients with SSNS and 2339 genetic ancestry matched controls obtained from the UK Biobank. We then performed a trans-ethnic meta-analysis with a previously reported European cohort of 422 pediatric patients and 5642 controls.

**Results:** Our GWAS confirmed the previously reported association of SSNS with *HLA-DR/DQ* (rs9271602, p=1.12×10^−27^, odds ratio[OR]=2.75). Trans-ethnic meta-analysis replicated these findings and identified a novel association at *AHI1* (rs2746432, p=2.79×10^−8^, OR=1.37), which was also replicated in an independent South Asian cohort. *AHI1* is implicated in ciliary protein transport and immune dysregulation, with rare variation in this gene contributing to Joubert syndrome type 3.

**Conclusions:** Common variation in *AHI1* confers risk of the development of SSNS in both Sri Lankan and European populations. The association with common variation in *AHI1* further supports the role of immune dysregulation in the pathogenesis of SSNS and demonstrates that variation across the allele frequency spectrum in a gene can contribute to disparate monogenic and polygenic diseases.

**AUTHOR SUMMARY:** Steroid-sensitive nephrotic syndrome (SSNS) is the most common kidney disease in children worldwide, but the cause of disease is not well understood. Genome-wide association studies (GWAS) in SSNS have shown that genes in the classical HLA region (the human immune centre) and several genes outside of this region are associated with the disease, which has allowed us to further understand the cause of disease. We performed a GWAS of Sri Lankan ancestry that included 420 paediatric patients and 2339 ancestry-matched controls and confirmed association at *HLA-DQ/DR* with SSNS. We then performed a Sri Lankan-European trans-ethnic meta-analysis and identified a new association with SSNS outside of HLA, in *AHI1*. This finding further supports the role of immune system involvement in the etiology of SSNS and increases our knowledge of the genetic causes of disease. *AHI1* is a gene that can also cause ciliary problems and demonstrates that different genetic variants within the same gene can contribute to both single-gene (Joubert syndrome, a rare disease that causes kidney and neurological problems) and multi-gene diseases (SSNS).

## INTRODUCTION

Steroid-sensitive nephrotic syndrome (SSNS) is the most common kidney disease in children worldwide, with an incidence of approximately 1-10 per 100,000(1). Incidence varies with ancestry; individuals of South Asian ancestry demonstrate higher risk for the disease compared with Europeans(2). These observations suggest genetic and/or environmental influences in the development of SSNS. Although there have been several studies that have identified genetic risk loci in children with SSNS of European and other ancestries(3–6), additional genetic risk factors associated with SSNS in children of South Asian ancestry have not been reported.

SSNS is characterized by the leakage of protein from the blood into the urine through damaged glomeruli(7). The etiology of SSNS remains unclear, although clinical observations suggest an underlying immunologic basis to disease: First, SSNS is defined by response to initial treatment with corticosteroid therapy and in patients that develop a relapsing course of disease, SSNS also responds to additional immunosuppressive medications(8). Second, the onset of disease is typically associated with a preceding infection, suggesting that prior activation of the immune system may trigger the disease(9). Third, antibodies directed toward Nephrin, a protein in the slit diaphragm in the glomerulus, were recently identified in patients with SSNS(10). These clinical observations suggest that SSNS is an autoimmune disorder, implicating both genetic and environmental factors contributing to development of the disease.

Genome-wide association studies (GWAS) have been instrumental in elucidating genetic risk factors for developing SSNS in childhood. The *HLA-DR/DQ* region has exhibited the strongest association with disease in European, South Asian, and Japanese populations(3–6,11), supporting the inference from clinical observations that SSNS has an immunological basis. Beyond the *HLA* region, genome-wide associations at *CALHM6* and *PARM1* have been identified in European children(3), and at *NPHS1* and *TNFSF15* in Japanese children(6). In the latter study, the *NPHS1* and *TNFSF15* loci were not replicated in a European population, suggesting that SSNS possesses different genetic architecture outside of *HLA* in these two different groups. We set out to perform a GWAS in a Sri Lankan population, followed by a European-Sri Lankan trans-ethnic meta-analysis, to identify additional genetic loci associated with SSNS to aid in further understanding of the pathophysiology of disease.

## RESULTS

### GWAS Study Cohort

A total of 663 individuals with childhood-onset SSNS and South Asian ancestry were available for our study, and 420 Sri Lankan cases were included in the association analysis following quality control. The control dataset was obtained from the UK Biobank from cohorts of self-reported Indian, Bangladeshi, Pakistani, Any other Asian background, and Other ethnic group ancestry for an initial total of 14,398 individuals. After ancestrally matching these individuals to our cases and performing quality control, 2339 healthy individuals with genetically determined Sri Lankan ancestry were included in the association analysis, with the majority obtained from the Any other Asian background and Other ethnic group cohorts. The case and control cohorts were imputed and combined to yield a total of 5,265,125 high quality SNPs for analysis.

### GWAS Results

GWAS of the Sri Lankan population showed two independent genome-wide significant signals (Figure 1, Table 1). The strongest association was detected in *HLA-DQA1* (rs9271602, p=1.12×10^−27^, OR=2.75, 95%CI 2.29-3.30) (Figure 2A). Conditional analysis on rs9271602 revealed a second independent signal at rs9391784 (p=1.16×10^−15^); further conditioning on rs9271602 and rs9391784 revealed a third signal at rs17212846 (p=2.57×10^−13^); and conditioning on rs9271602, rs9391784, and rs17212846 revealed a fourth signal at rs9260172 (p=3.27×10^−9^)(see Supplementary Figure S4).

**Figure 1:**
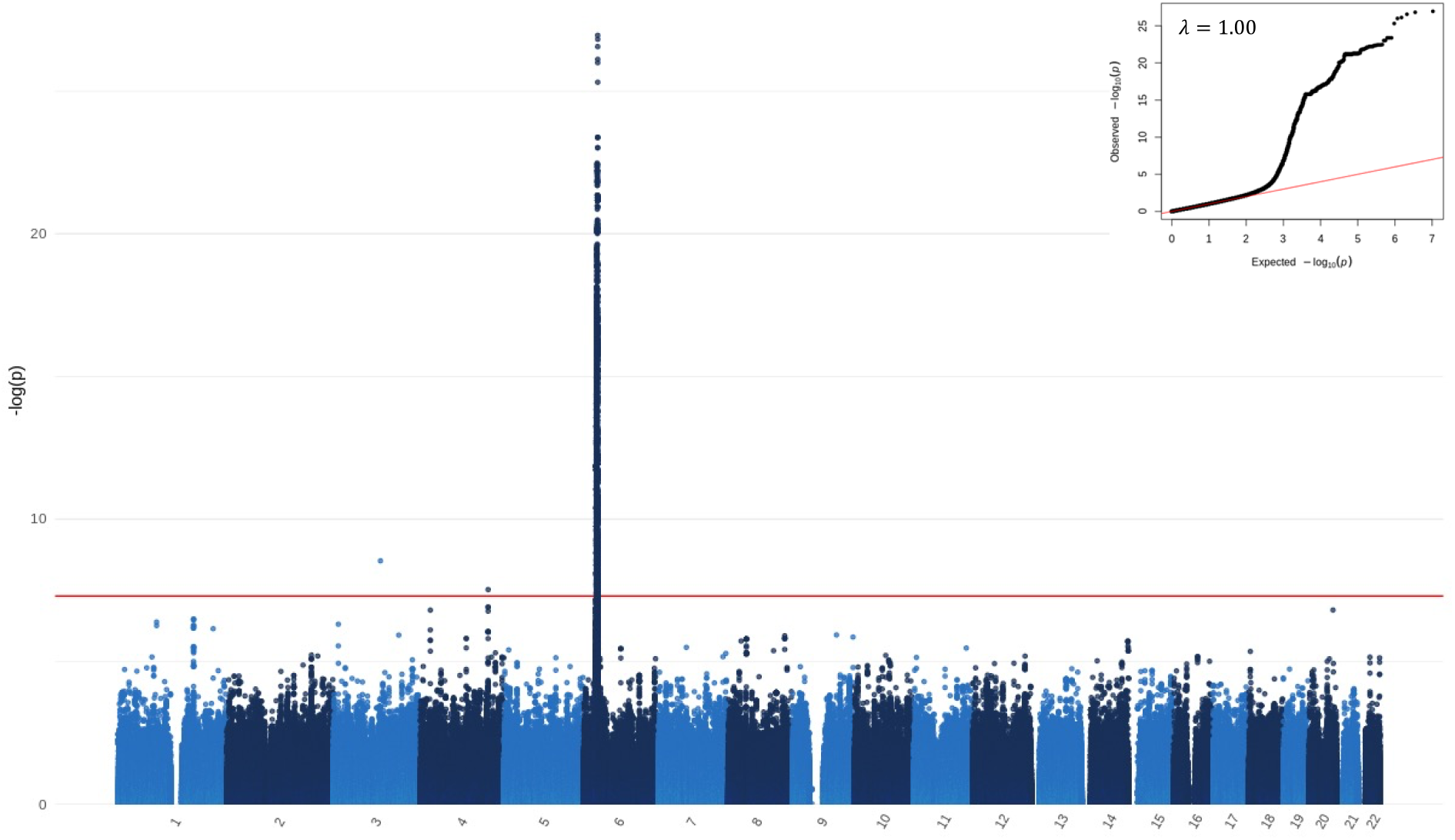
Manhattan plot in Sri Lankan SSNS. Genome-wide association study (GWAS) for steroid sensitive nephrotic syndrome (SSNS) in 420 Sri Lankan patients and 2339 ancestrally matched controls. Mixed model logistic regression analysis adjusted for the first three principal components was performed in SAIGE. Autosomal chromosomes (1-22) are listed along the x axis. The level of significance is depicted along the y axis as -log_10_(p). Each dot represents a variant. The red line represents the threshold of genome-wide significance (p=5×10^−8^). Three loci achieve genome wide significance on chromosomes 3, 4, and 6. QQ-plot and lambda are displayed in the top right corner.

**Table 1:** Lead SNPs associated with Sri Lankan SSNS. SNP, single nucleotide polymorphism; DR^2^, imputation dosage R^2^; AF, allele frequency; OR, odds ratio; CI, confidence interval.

| SNP | Locus | Gene | DR<sup>2</sup> | Test Allele | AF Cases | AF controls | OR | 95% CI | p-value |
| --- | --- | --- | --- | --- | --- | --- | --- | --- | --- |
| rs9271602 | 6p21.32 | *HLA-DQA1* | 1 | G | 0.53 | 0.29 | 2.75 | 2.29-3.30 | $1.12 \times 10^{-27}$ |
| rs78120384 | 3q13.12 | Intergenic | 1 | A | 0.18 | 0.09 | 2.34 | 1.76-3.09 | $2.91 \times 10^{-9}$ |
| rs74537360 | 4q31.3 | *TMEM131L* | 1 | T | 0.14 | 0.07 | 2.37 | 1.75-3.22 | $2.98 \times 10^{-8}$ |

**Figure 2:**
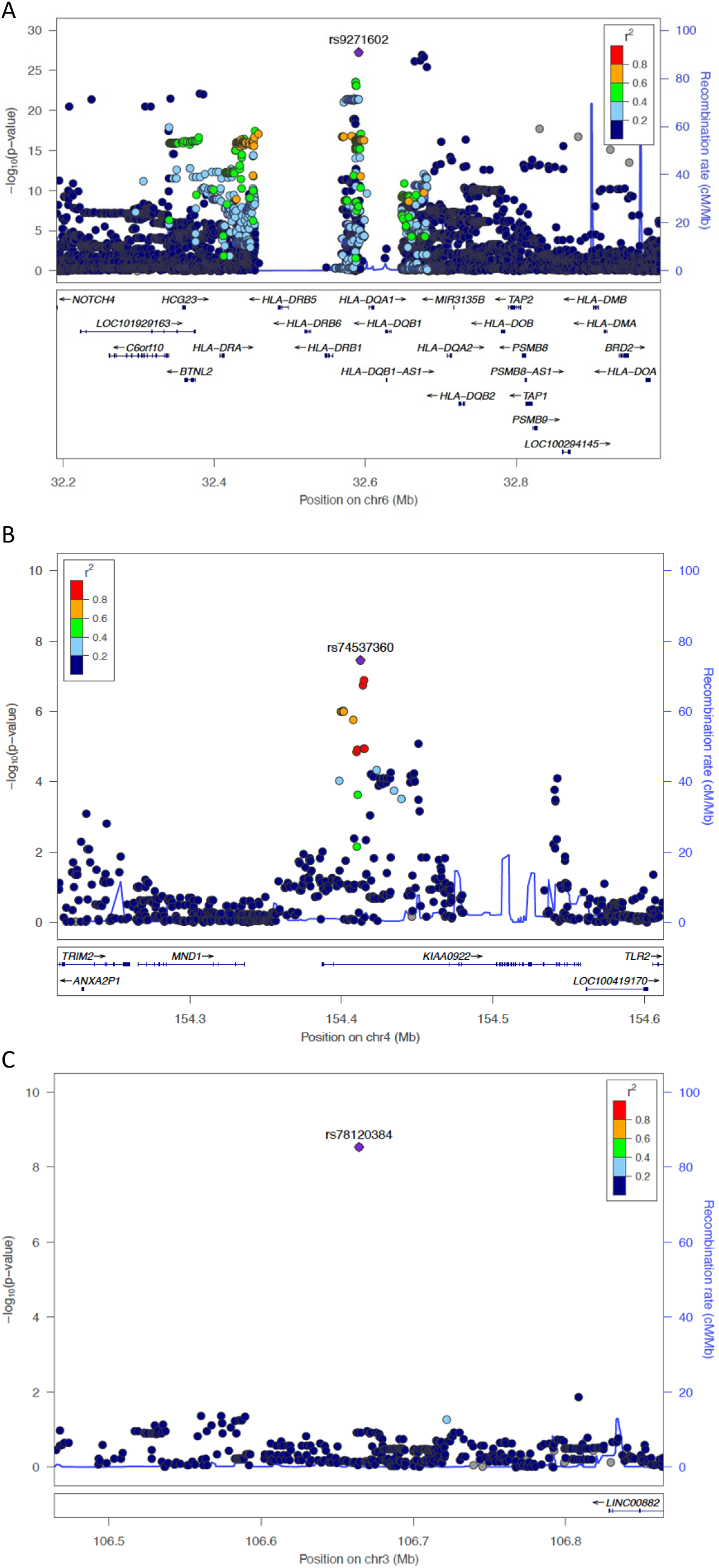
Locus zoom plot for regions on chromosomes 6p21.32 and 4q31.3 identified in Sri Lankan discovery cohort. Index SNPs are annotated as a purple diamond over the respective genes, listed below. The surrounding SNPs coloured in yellow and green are in linkage disequilibrium with the index SNP as depicted by the r^2^ value in the legend. Genes and positions in megabases (Mb) are listed along the x axis. The level of significance is depicted along the y axis as -log_10_(p). Each dot represents a variant. A) Lead SNP (rs9271602) in *HLA-DQ/DR* region; B) lead SNP (rs74537360) in the *KIAA0922* (otherwise known as *TMEM131L*) gene; and C) lead SNP (rs78120384), upstream of the gene *LINC00882*. No other SNPs in LD with rs78120384 were associated with disease, suggesting a false positive result.

The lead SNP at *HLA-DQA1* is in strong LD with rs2858317 and rs3828799 (identified by Dufek *et al*)(3), rs4642516 (identified by Jia *et al*)(4), rs1129740 and rs1071630 (identified by Gbadegesin *et al)*(11), and rs1063348 and rs28366266 (identified by Debiec *et al*)(5).

The next strongest association was outside the HLA region at 4q31.3 in the gene, *TMEM131L*, previously called *KIAA0922* (rs74537360, p=2.98×10^−8^, OR=2.37, 95%CI 1.75-3.22) (Figure 2B). Genome-wide significance was lost after conditioning on rs74537360 (Supplementary Figure S5). A further isolated marker (rs78120384) outside of HLA reached genome-wide significance, which was on the lower border of accepted allele frequencies and was deemed a false positive result (Figure 2C).

The power of this GWAS exceeded 80% to detect common alleles (MAF >0.01) with genotypic relative risk (RR) > 2.2 at a significance threshold of p>5×10^−8^ under an additive model. The inflation factor (λ) was calculated to be 1.00 suggesting no evidence of genomic inflation.

### HLA Fine-mapping

Significant association with SSNS was detected in six *HLA* alleles, including three previously reported subtypes associated with SSNS in Europeans: *DQB1*02:01, DQA1*01*, and *DQA1*02:01*(3) (Table 2). The strongest association was observed in *DQB1*02:01*, which was a risk haplotype. The strongest protective allele was in *HLA-DQA1*01*. Conditional analysis on the lead HLA allele, *HLA-DQB1*02:01*, revealed that the only further independent signal was in *HLA-B*52:01* (Supplementary Figure S6).

**Table 2:**
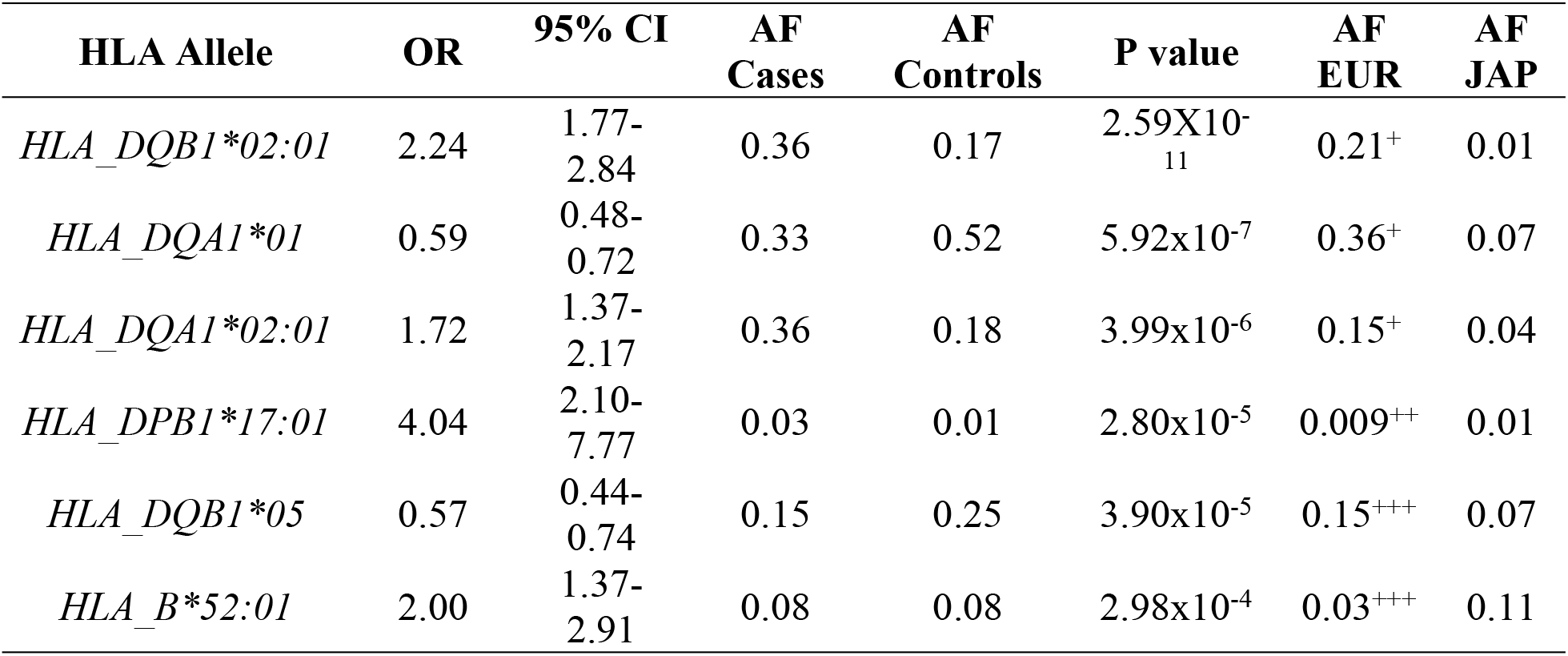
Classical HLA alleles associated with Sri Lankan SSNS. OR, Odds Ratio; CI, confidence interval; AF, allele frequency; AF, allele frequency; EUR, European; JAP, Japanese. ^+^ values obtained from Dufek *et al(3)*; ^++^ values obtained from Lemin *et al(12)*; ^+++^ values obtained from Tokic *et al(13)*; all allele frequency for the Japanese population were obtained from http://hla.or.jp

### Trans-ethnic meta-analysis

Trans-ethnic meta-analysis was performed with the previously reported European GWAS by Dufek *et al*(3), and showed replication of the *HLA-DQ/DR* locus (rs2856665, p=2.45×10^−68^, OR=4.06, 95%CI 3.47-4.75). The European-identified signal in 6q22.1 (*CALHM6*) was also replicated, though was primarily driven by the European cohort (rs2637681, p=6.69 x10^−13^, OR 0.62, 95%CI 0.54-0.71). A novel association, driven by a combination of both cohorts, was identified at 6q23.3 in the gene *AHI1* (rs2746432, p=2.79×10^−8^, OR 1.37, 95% CI 1.22-1.52). See Table 3, Supplementary Table S1, and Figures 3 and 4. The signal in *TMEM131L* was not replicated, though the set of overlapping markers used in the meta-analysis did not include the lead SNP at this locus.

**Table 3:**
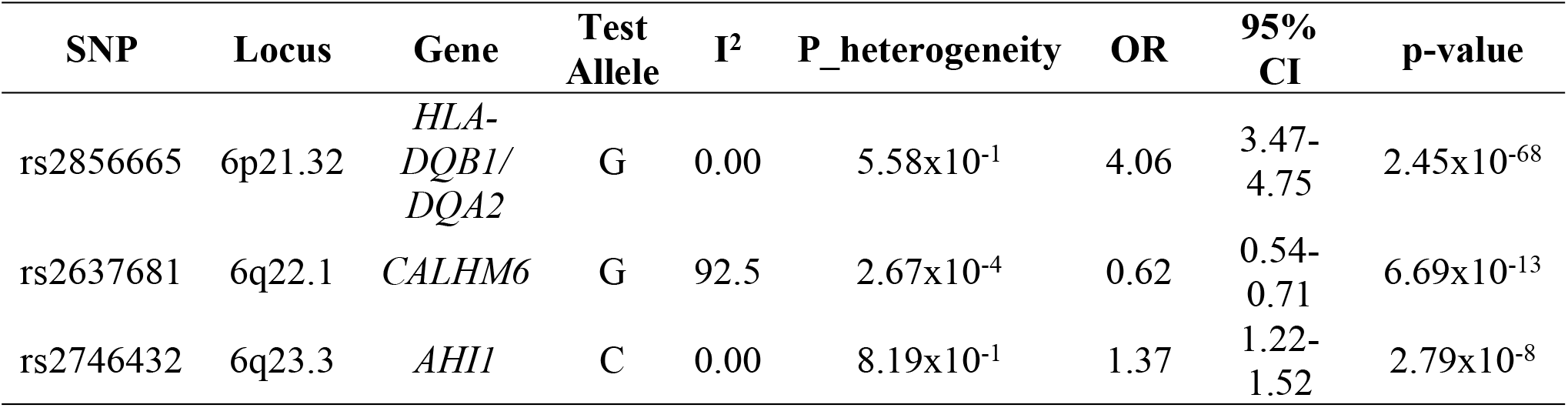
Lead SNPs associated with SSNS in Sri Lankan-European meta-analysis of childhood SSNS. SNP, single nucleotide polymorphism; I^2^, percentage of total variation across studies due to heterogeneity; P_heterogeneity, p-value of Cochrane Q test for heterogeneity; CI, confidence interval

**Figure 3:**
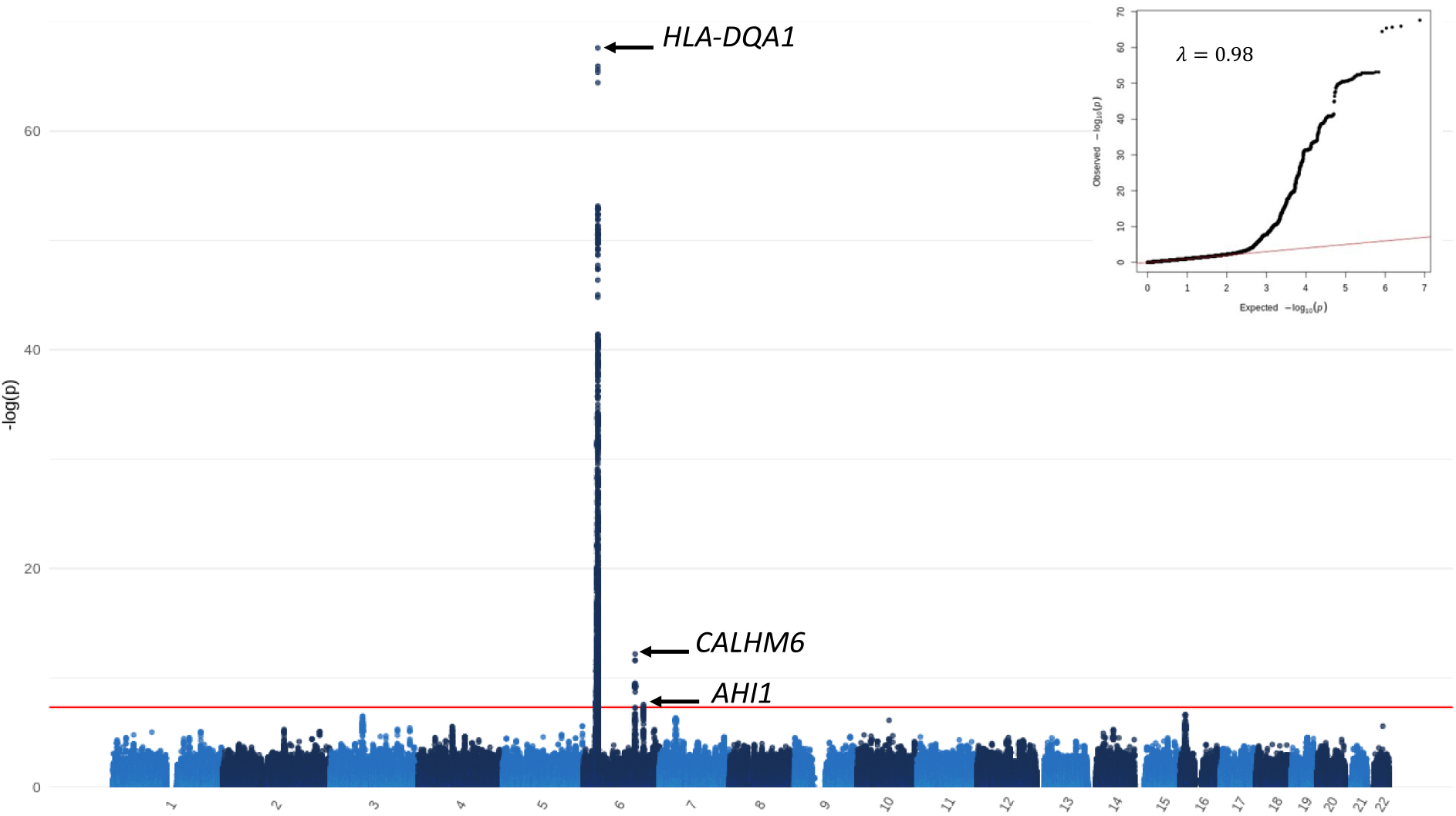
Manhattan plot of the trans-ethnic meta-analysis of the Sri Lankan discovery cohort and the European replication cohort. Autosomal chromosomes (1-22) are listed along the x axis. The level of significance is depicted along the y axis as -log_10_(p). Each dot represents a variant. The red line represents the threshold of genome-wide significance (p=5×10^−8^). The inverse-variance method based on a fixed-effects model was used. Three loci achieve genome wide significance on chromosome 6; variants in *HLA-DQA1, CALHM6*, and *AHI1* are labelled on the plot. QQ-plot and lambda are displayed in the top right corner.

**Figure 4:**
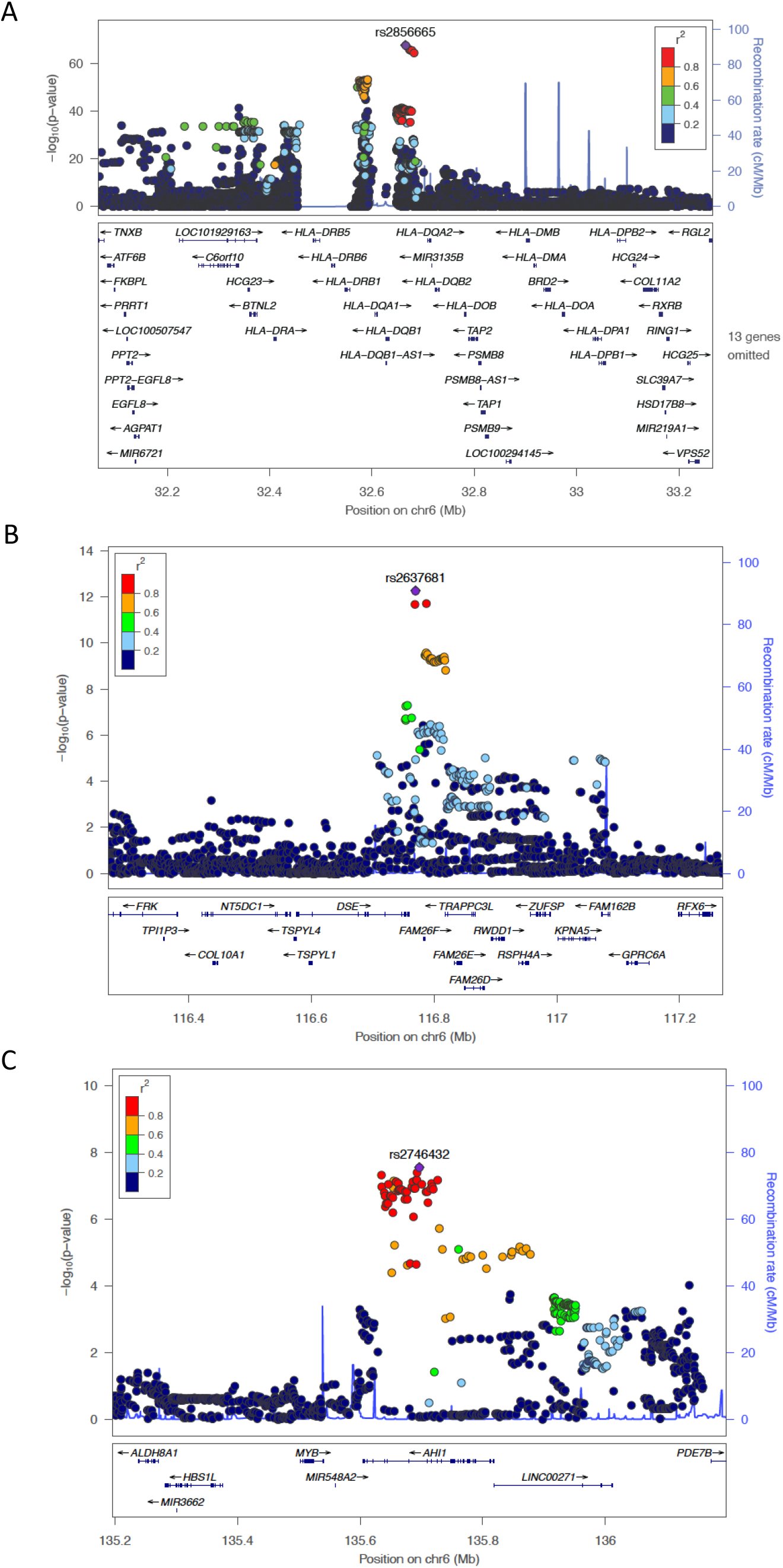
Locus zoom plot for regions on chromosomes 6p21.32, 6q22.1, and 6q23.3 in trans-ethnic meta-analysis. Index SNPs are annotated as a purple diamond over the respective genes, listed below. The surrounding SNPs coloured in yellow and green are in linkage disequilibrium with the index SNP as depicted by the r^2^ value in the legend. Genes and positions in megabases (Mb) are listed along the x axis. The level of significance is depicted along the y axis as -log_10_(p). Each dot represents a variant. A) Lead SNP (rs2856665) in *HLA-DQ/DR* region; B) lead SNP (rs2637681) in the *FAM26F* (otherwise known as *CALHM6*) gene; and C) lead SNP (rs2746432) in the *AHI1* gene.

### Replication

The novel genome-wide significant signal in *AHI1* was replicated in an independent South Asian population (INSIGHT cohort) (rs2746432, p=1.13×10^−2^, OR 1.58), although the power to do so was only 0.466 (Table 4). This lead SNP also showed evidence of association in the Japanese cohort published by Jia *et al* (rs2746432, p=1.08×10^−3^)(6). The signal in *TMEM131L* was not replicated in this cohort (rs74537360, p=0.76, OR 1.09), despite power to detect this signal being 0.894.

**Table 4:**
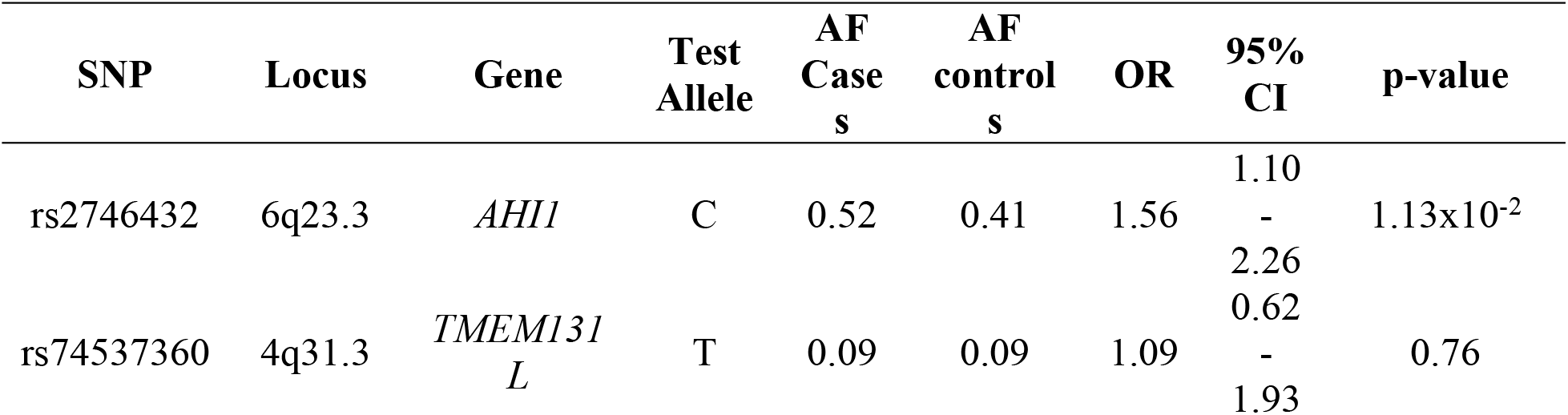
Replication of candidate SNPs associated with Sri Lankan SSNS in INSIGHT cohort. SNP, single nucleotide polymorphism; AF, allele frequency; OR, odds ratio; CI, confidence interval.

### Gene annotation

The lead SNP at the *AHI1* locus (rs2746432) exhibits cis-eQTLs in the GTEx(14) database in multiple tissues. Notably, rs2746432 shows strong cis-eQTL effects in fibroblasts (NES 0.51, p=1.6×10^−28^), EBV-transformed lymphocytes (NES 0.60, p=2.6×10^−9^), and in the spleen (NES 0.41, p=6.0×10^−9^). The SSNS risk (minor) allele in rs2746432 decreased the expression of *AHI1* in all cell types, indicating that in cases where the risk allele was more frequent, the expression of *AHI1* is downregulated.

## DISCUSSION

The present Sri Lankan GWAS and trans-ethnic meta-analyses were performed in the largest South Asian cohort to date and identified common variants in *AHI1* as a new susceptibility locus for childhood SSNS. This study has also confirmed previous association findings of SSNS with *HLA-DQ/DR*. Furthermore, the larger sample size enabled additional fine mapping of the *HLA* locus. These findings provide new insights into our understanding of the genetic background of childhood SSNS, and further support an immunologic basis to its pathogenesis. This study also provides evidence of association with alleles at *TMEM131L*, but this was not replicated in either the meta-analysis or in an independent South Asian cohort, suggesting that this association is either a Type 1 error or represents a genetic risk factor uniquely found in the Sri Lankan (as opposed to the South Asian or European) population. These findings suggest that the higher prevalence of SSNS in Sri Lankan compared to European individuals is not explained by common genetic variants (within the limitations of this study’s power to detect association).

### HLA Locus

The strongest association in our analysis was in the HLA region. In the Sri Lankan discovery cohort, the lead SNP, rs9271602, was in the *HLA-DR/DQ* region, specifically in the *HLA-DQA1* and *HLA-DQB1* genes. This finding was replicated in the trans-ethnic meta-analysis of the European and Sri Lankan cohorts, with the strongest association at rs2856665, between *HLA-DQB1* and *HLA-DQA2* but with linkage disequilibrium extending to *HLA-DQA1*. All previous GWAS published on SSNS have found association within these genes, including populations of European, South Asian, and Japanese ancestry(3–6,15).

Fine-mapping of the HLA alleles identified *HLA-DQB1*02:01, HLA-DQA1*02:01, HLA-DPB1*17:01*, and *HLA-B*52:01* to be associated with increased risk of SSNS. Of these, *HLA-DQA1*02:01* and *HLA-DQB1*02* were also found to be associated with increased risk of disease in European and South Asian studies(3,5,15). *HLA-DQA1*01* and *HLA-DQB1*05* were the protective alleles associated with SSNS in our Sri Lankan discovery cohort, with the *HLA-DQA1*01* allele replicating in European(3) and South Asian(15) populations. Conversely, in Japanese populations, altogether different risk and protective HLA alleles have been identified(4,6). These findings demonstrate substantial overlap between European and South Asian populations, but not Japanese. This is most likely explained by differing allele frequencies in the different populations (see Table 2 and Supplementary Table S2).

In the conditional analysis on the lead allele, *HLA-DQB1*02:01*, in our Sri Lankan discovery cohort, both the risk and protective alleles at *HLA-DR/DQ* disappeared, leaving only *HLA-B*52:01* as the independent signal. *HLA-B*52:01* (the most common subtype of allele at B*52) was the only class I HLA allele associated with disease in our discovery GWAS analysis. Though this allele was relatively rare in the Sri Lankan population (MAF 0.08), it is interesting because of its association with several other immune-mediated diseases, including ulcerative colitis(16) and Takayasu’s arteritis(17).

### Association at *AHI1*

The novel signal outside of HLA detected in our trans-ethnic meta-analysis was on chromosome 6 within the gene *AHI1 (Abelson Helper Integration Site 1)* (rs2746432, p=2.80×10^−8^, OR 1.36, 95% CI 1.22-1.52)(18). *AHI1* encodes the protein, Jouberin, which is a component of a ring-like protein complex in the transition zone at the base of cilia. Together, with the other proteins that compose the complex, *AHI1* acts to restrict protein diffusion between the plasma and ciliary membranes; disruption of the complex leads to reduction in cilia formation and a reduction in signalling receptors from the remaining cilia(19). Rare biallelic mutations in *AHI1* cause Joubert syndrome, a rare monogenic disorder manifesting in agenesis of the cerebellum, ataxia, hypotonia, and intellectual disabilities(20). Interestingly, our meta-analysis revealed that common variants in *AHI1* are associated with SSNS.

*AHI1* has a diverse array of biological functions. It is known to be important in the kidney through its interaction with *NPHP1 (nephrocystin-1)*, which encodes another protein at the basal body of cilia. Mutations in *NPHP1* are associated with Joubert syndrome accompanied by renal dysfunction, accounting for the majority of cases of nephronophthisis(21). *AHI1* and *NPHP1* form heterodimers and heterotetramers, and mutations in *AHI1* have been shown to change this binding pattern(20).

*TTC21B*, a ciliary gene, has recently been found to be associated with familial FSGS(22). The gene product of *TTC21B*, IFT139, was shown to localise at the base of the primary cilium in developing podocytes from fetal kidney tissue, and IFT139-deficient podocytes were shown to have primary ciliary defects, abnormal cell migration, and alterations in the cytoskeleton. While these findings demonstrate the importance of a ciliary gene in glomerular function and disease(22), a precise mechanism linking variation at *AHI1* causing ciliary dysfunction and SSNS remains to be elucidated.

*AHI1* is also involved in immune system function. Jiang *et al* found that *AHI1* is highly expressed in primitive types of normal hematopoietic cells and is downregulated during early differentiation(23). Thus, alterations in *AHI1* expression may contribute to the development of certain types of human leukemias. Notably, a GWAS in the autoimmune disease multiple sclerosis (MS) detected a susceptibility variant in *AHI1* (rs4896153) that was subsequently shown to have strong *cis-*eQTL effect on overall *AHI1* expression(24). Functional studies showed that expression peaked after stimulation of human CD4+ T cells, suggesting that it may play a role in early T cell receptor (TCR) activation. *AHI1* has also been shown to be involved in actin organization(25), and therefore the authors of this study speculated that *AHI1* may play a role in the formation or stabilization of the TCR synapse as a mechanism for its association with MS(24).

The eQTL analysis of the lead SNP, rs2746432, showed *cis*-eQTL effects on *AHI1* in EBV-transformed lymphocytes, with the risk allele at this variant associated with decreased *AHI1* expression. Thus, it is possible that decreased *AHI1* expression in the lymphocytes of individuals with SSNS could lead to increased cytokine production, and/or destabilization of the T cell receptor complex, both resulting in immune-system dysregulation. The association of common variation at *AHI1* with SSNS in addition to the established association of (biallelic) rare variants of *AHI1* with Joubert syndrome demonstrates that variation across the allele frequency spectrum in a gene can contribute to both monogenic and polygenic disease, and that these alleles might act by different mechanisms, resulting in altogether different disorders.

### Association at *TMEM131L*

The strongest signal outside of HLA in our Sri Lankan discovery GWAS was on chromosome 4 at 4q31.3 (rs74537360, p=3.47×10^−8^, OR=2.36, 95%CI 1.74-3.20), which was not replicated in our meta-analysis or in the independent South Asian cohort. However, this SNP (rs74537360) was not present in our meta-analysis (due to its absence in the European cohort), and therefore a proxy SNP (rs4696465, R^2^=0.87) was used to determine replication (see Supplementary Table S1). The lead SNP is located in the gene *KIAA0922*, otherwise known as *TMEM131L (Transmembrane 131 Like)*(26). *TMEM131L* is a transmembrane protein that has been shown to down regulate immature T cell proliferation in the thymus, and to inhibit Wnt signalling(27), which is an important pathway for immune cell maintenance and renewal. *TMEM131L* has also been shown to inhibit the Notch signalling pathway, which is a highly conserved signalling pathway that is crucial in the malignant transformation of cells(28). Furthermore, expression data shows that *TMEM131L* is highly expressed in both B and T lymphocytes. Taken together, these observations suggest that *TMEM131L* is a plausible candidate gene in the susceptibility to SSNS, consistent with abundance of clinical evidence that SSNS is an immune-mediated disease. A variant in the *TMEM131L/TLR2* region (not in linkage disequilibrium with our lead SNP) was found to be associated with another autoimmune disease, systemic lupus erythematosus (SLE)(29), providing additional evidence for the importance of *TMEM131L* in immune regulation. However, the absence of replication of our initial signal in a second South Asian cohort despite adequate power to do so (and similar frequency of the relevant allele in numerous populations, see Supplementary Table S1), suggests that, while this could represent a uniquely Sri Lankan genetic risk factor for the disease, the possibility that this apparent association represents a Type 1 error remains, so (pending further independent data confirming this association) it should be regarded as a preliminary finding.

### Association at previously published non-HLA loci

#### CALHM6

The strongest association outside of HLA in our trans-ethnic meta-analysis was on chromosome 6 (rs2637681, p=5.44 x10^−13^, OR 0.62, 95%CI 0.54-0.70). This marker is in the gene, *CALHM6 (Calcium Homeostasis Modulator Family Member 6)*(30). This locus was associated with SSNS in the previously published European GWAS(3), and was also reported as a potential signal in the SSNS GWAS by Debiec *et al*(31). However, it was not significantly associated with disease in our Sri Lankan discovery cohort (see Supplementary Table S1). Unsurprisingly, association of rs2637681 in our trans-ethnic meta-analysis was mainly driven by the European cohort. The direction of effect was the same in both cohorts, however, which supports the relevance of this finding.

Power calculation indicated that the Sri Lankan discovery GWAS was powered to detect a signal of similar strength at this locus at p<0.05, even considering the lower frequency of the associated allele in the Sri Lankan population (0.162 compared with 0.413 in Europeans). However, the observed signal (p=0.199, OR 0.86, 95%CI 0.69-1.08) was not as strong as this. There are several potential explanations for this, including differences in LD patterns in individuals of different ancestries, or that there is a true difference in effect size at this variant, perhaps due to differences in genetic background or environmental exposures(32). It has been previously demonstrated that variants associated with a particular disease in one ancestral group are not always reproduced in another(33). Furthermore, variants shared between autoimmune disorders have been shown to be protective in one disorder and risky in another(34).

#### PARM1

The lead SNP (rs10518133) at the *PARM1* locus on chromosome 4 from the largest and most recent European GWAS was replicated in our analysis (p=2.90×10^−6^, OR 1.59, 95%CI 1.31-1.93, see Supplementary Table S1)(3). However, the signal was entirely driven by the European cohort: p=2.50×10^−8^, OR 1.96, 95% CI 1.57-2.45 in the European cohort, which was not seen in the Sri Lankan cohort (p=0.403, OR 0.85, 95% CI 0.57-1.25). As was the case for the *CALHM6* locus, our Sri Lankan discovery cohort was sufficiently powered to replicate this finding, and we believe the lack of replication is best explained by differences in either LD pattern or effect on disease risk of alleles at this locus in the two populations.

#### NPHS1/KIRREL2

The lead variant at *NPHS1/KIRREL2* (rs56117924) identified by Jia *et al*(6) in the Japanese population was not captured in our analysis. A suitable proxy marker in high LD with rs56117924 was also not available (highest R^2^=0.04 in meta-analysis). Therefore, replication at this locus was not possible in our study. Interestingly, the allele frequency of the test allele (A) at rs56117924 in European populations is zero (*i*.*e*. 100% of Europeans have the genotype GG at this marker), which is the likely explanation for why this SNP and haploblock were not captured in our dataset(35).

#### TNFSF15

Jia *et al*(6) also identified *TNFSF15* (lead SNP rs6478109) to be associated with childhood SSNS in the Japanese population. This signal was replicated in our meta-analysis (p=9.96×10^−3^, OR 0.85, 95%CI 0.75-0.96, see Supplementary Table S1), and the direction of effect was the same (protective) in both the Sri Lankan and European cohorts. In the original Japanese study, the authors replicated this locus in an independent South Asian cohort, but not in a European cohort(6). Based on the results of our study, *TNFSF15*, in addition to *AHI1*, are therefore the first non-HLA loci associated with SSNS in all three populations studied (South Asian, Japanese, and European).

### Limitations

This study has several limitations. Firstly, there was limited clinical information on the individuals included in the study. Details such as age of onset or relapse pattern could have enhanced our understanding of the relationship between markers of clinical severity and number of risk alleles, although the relatively small size of the cohort would limit the power to perform this type of analysis. Secondly, the case and control datasets were genotyped on different platform, which limited the number of overlapping markers. We overcame this problem by filtering for genotyping discrepancies and imputing the datasets separately, but this has greater potential for error than if the case and control datasets were genotyped on the same platform.

## CONCLUSIONS

In summary, our study showed a novel association of childhood SSNS with alleles at *AHI1* and confirmed previous associations at *HLA-DR/DQ*. These findings further support the role of immune dysregulation in the pathophysiology of disease. The *AHI1* association, in particular, suggests a link between a ciliary gene and glomerular disease and reinforces an emerging paradigm in nephrology: in genes harbouring rare Mendelian variants, common alleles can increase the susceptibility of polygenic diseases. Our study also illustrates the importance of performing GWAS in diverse populations, because by doing so, we were able to increase the power to detect a novel variant associated with SSNS.

## MATERIALS AND METHODS

Abbreviated methods follow. Detailed methods may be found in the Supplementary Material.

### Study Populations

Sri Lankan patients diagnosed with childhood SSNS (age of onset <18 years) were recruited into the study. Most patients were of self-reported Sri Lankan ancestry, with additional ancestrally matched patients identified by principal component analysis (PCA). All patients were diagnosed with SSNS as per the KDIGO guidelines(36). Patients were recruited by collaborating clinicians at their affiliated institutions, as well as from the Prednisolone in Nephrotic Syndrome (PREDNOS, EudraCT 2010-022489-29) and PREDNOS2 (EudraCT 2012-003476-39) trials(37,38). Informed verbal and/or written consent was obtained from each participant and ethical approval was granted at each contributing institution. Ancestrally matched controls were obtained from the UK Biobank(39).

### Genotyping, quality control, and whole-genome imputation

Isolation of DNA, genotyping, quality control, PCA, and imputation were performed using standard procedures (See Supplementary Methods and Supplementary Figures S1-S3). Patients were genotyped *via* the Infinium Multi-Ethnic Global Array (MEGA) BeadChip v.A1 at University College London Genomics (Institute of Child Health, University College London, UK). UK Biobank controls had been genotyped using the Applied Biosystems UK Biobank Axiom Array.

Whole-genome imputation was performed with minimac4 on the Michigan Imputation Server (MIS)(40,41) using the 1000 Genomes Project Phase 3 as the reference panel (42). PLINK versions 1.90 and 2.00 were used for quality control analysis(43).

### Genome-wide association analysis

Genome-wide association analysis was performed in SAIGE(44) with adjustment for sex and the first three principal components of ancestry. Using more than three principal components resulted in genomic deflation, suggesting over-fitting. Conditional analysis of the lead SNPs was performed in SAIGE using the same model adjusted for sex and principal components. A genome-wide significance threshold of p<5×10^−8^ was used. R v4.2.1 was used to generate Manhattan plots. Regional plots were generated using LocusZoom with 1000 Genomes Nov 2014 used as the linkage disequilibrium (LD) reference(45).

### HLA Fine-Mapping

HLA imputation was performed with minimac4 on the Michigan Imputation Server using the HLA-TAPAS (HLA-Typing At Protein for Association Studies) reference panel(40,41,46). HLA association analysis was performed in PLINK v2.00 using a logistic regression model adjusted for sex and the first ten principal components of ancestry. Conditional analysis of the lead HLA allele was performed using the same logistic regression model adjusted for sex and principal components. A significance threshold of p<3×10^−4^ (0.05/136) was used to adjust for multiple comparisons with the n=136 4-digit HLA alleles used in the analysis.

### Trans-ethnic meta-analysis

Trans-ethnic meta-analysis was performed with a previously reported GWAS of European children with SSNS(3). Analysis was conducted using the set of overlapping markers between the two datasets. The inverse-variance method was used based on a fixed-effects model in META: https://mathgen.stats.ox.ac.uk/genetics_software/meta/meta.html. The genomic inflation factor (λ) and population sizes of each study were incorporated into the model. Results were considered significantly heterogeneous with a Cochran Q test p-value <0.10. The genome-wide significance threshold for the meta-analysis was considered p<5×10^−8^.

### Replication

Replication of the two novel candidate SNPs (in *TMEM131L* and *AHI1*) was assessed in an independent population that comprised 150 South Asian (including Sri Lankan) participants from the INSIGHT(47) cohort and 277 controls from the Spit for Science study(48). South Asian genetic ancestry was determined by principal component analysis using 1000 Genomes(42) ancestry controls as reference. Association analyses were carried out under an additive model. Significance threshold for replication was considered as p<0.05/2=0.025. For replication of candidate SNPs at the *HLA* locus, we examined the results of previously published GWAS in SSNS(3–6).

### Power calculation

The GWAS and replication study power were calculated using the Michigan Genetic Association Study power calculator(49) assuming a disease prevalence of 1:10,000. For the initial GWAS (420 cases and 2339 controls), the minimum genotype relative risk with a power of 0.8 was calculated using an additive model assuming a disease allele frequency of 0.10 in the control population and a significance level of 5×10^−8^. For the replication analysis (150 cases and 277 controls), power calculation assumed the genotype relative risk and allele frequency at each locus observed in the discovery GWAS, with a significance threshold of p<0.025.

## ACKNOWLEDGEMENTS

We would like to acknowledge the patients and families who participated in this research, without direct benefit to themselves.

## CONFLICT OF INTEREST STATEMENT

Nothing to disclose.

The results presented in this paper have not been published previously in whole or part.

## AUTHOR CONTRIBUTIONS

M.L. Downie designed the study, performed quality control and association analysis, interpreted results, drafted the manuscript, and approved of the manuscript as written.

S. Gupta contributed to data collection, interpreted results, critically revised the manuscript, and approved of the manuscript as written.

C. Voinescu provided computing support, interpreted results, critically revised the manuscript, and approved of the manuscript as written.

A.P. Levine designed in-house computing software, critically revied the manuscript, and approved of the manuscript as written.

O. Sadeghi-Alavijeh provided computing support, interpreted results, critically revised the manuscript, and approved of the manuscript as written.

S. Dufek-Kamperis contributed to data collection, critically revised the manuscript, and approved of the manuscript as written.

J. Cao analyzed validation cohort data, critically revised the manuscript, and approved of the manuscript as written.

M. Christian contributed patient data and approved of the manuscript as written.

J.A. Kari contributed patient data and approved of the manuscript as written.

S. Thalgahagoda contributed patient data and approved of the manuscript as written.

R. Ranawaka contributed patient data and approved of the manuscript as written.

A. Abeyagunawardena contributed patient data and approved of the manuscript as written.

R. Gbadegesin contributed patient data and approved of the manuscript as written.

R. Parekh contributed patients data, interpreted results, critically revised the manuscript, and approved of the manuscript as written.

R. Kleta designed the study, interpreted results, critically revised the manuscript, and approved of the manuscript as written.

D. Bockenhauer designed the study, interpreted results, critically revised the manuscript, and approved of the manuscript as written.

H.C. Stanescu designed the study, interpreted results, critically revised the manuscript, and approved of the manuscript as written.

D.P. Gale designed the study, interpreted results, critically revised the manuscript, and approved of the manuscript as written.

## FUNDING

M.L. Downie and D.P. Gale are supported by St. Peter’s Trust for Kidney, Bladder, & Prostate Research. M.L. Downie was supported by the KRESCENT Post-Doctoral Fellowship from the Kidney Foundation of Canada.

## DATA AVAILABILITY STATEMENT

The data underlying this article will be shared via the European Genome-phenome Archive (EGA). Accession numbers and/or DOIs will be made available after acceptance.

## SUPPORTING INFORMATION CAPTIONS

Supplementary Methods

Supplementary Figure S1: Flowchart of the data processing for the SSNS Sri Lankan discovery GWAS

Supplementary Figure S2: Genotyping discrepancy analysis in Sri Lankan controls genotyped alongside Sri Lankan SSNS cases versus UK Biobank controls

Supplementary Figure S3: Principal component analysis of case-control dataset anchored by 100 Genomes controls

Supplementary Figure S4: Conditional analysis in HLA-DQ/DR region identified in Sri Lankan discovery cohort

Supplementary Figure S5: Conditional analysis in 4q31.3 region (*TMEM131L)* identified in Sri Lankan discovery cohort

Supplementary Figure S6: HLA 4-digit allele analysis in the Sri Lankan discovery cohort

Supplementary Table S1: Genome-wide significant variants associated with SSNS published in European, Japanese, and Sri Lankan populations

Supplementary Table S2: HLA alleles associated with SSNS in Sri Lankan and European populations

Supplementary References

## Notes

### Competing Interest Statement

The authors have declared no competing interest.

